# Regulatory Roles of Long Non-Coding RNAs in Arterial Stiffness and Hypertension: Insights from Two African American Studies

**DOI:** 10.1101/2024.08.11.607492

**Authors:** Jose D. Vargas, Malak Abbas, Gabriel Goodney, Amadou Gaye

## Abstract

**Background:** Arterial stiffness, commonly assessed via pulse wave velocity (PWV), is marked by reduced arterial elasticity and serves as a significant risk factor for cardiovascular disease and an early indicator of hypertension. This study investigated the regulatory roles of long non-coding RNAs (lncRNAs) in modulating mRNAs associated with arterial stiffness and hypertension, with a particular focus on African Americans, a population disproportionately impacted by hypertension.

**Methods:** We utilized whole-blood transcriptome sequencing data from two African American (AA) cohorts with high hypertension prevalence: the GENE-FORECAST study (436 subjects) and the MH-GRID study (179 subjects). Our objectives were to: (1) identify lncRNAs and mRNAs differentially expressed (DE) between the upper and lower tertiles of PWV, (2) determine DE lncRNAs associated with the expression levels of each DE mRNA, and (3) link the lncRNA-modulated mRNAs to hypertension across both datasets.

**Results:** Differential expression analysis revealed 1,035 DE mRNAs and 31 DE lncRNAs between upper and lower PWV groups. Then lncRNA-mRNA pairs significantly associated were identified, involving 31 unique lncRNAs and 1,034 unique mRNAs. Finally, 22 of the lncRNA-modulated mRNAs initially linked to PWV were found associated with hypertension, in both datasets. Interestingly, 30 lncRNAs were linked to the expression of UCP2 (Uncoupling Protein 2), a gene implicated in oxidative stress and endothelial function.

**Conclusions:** Our findings underscore the significant roles of lncRNAs in regulating gene expression associated with arterial stiffness and hypertension. The differential expression of UCP2 in relation to PWV and hypertension, along with its potential regulation by lncRNAs, offers valuable insights into the molecular mechanisms underlying arterial stiffness and its connection with hypertension.

## INTRODUCTION

Hypertension (HTN) is an important preventable risk factor for cardiovascular disease (CVD)^1^. As of 2010, 1.38 billion people were estimated to have hypertension (HTN) with its prevalence increasing due to increasing age and exposure to risk factors^2^. Racial/ethnic minorities have a higher burden of HTN with black adults in the US having the highest prevalence of HTN compared to other racial/ethnic groups^3^. Pulse wave velocity (PWV), an increasingly recognized measure of arterial stiffness, has been shown to accurately predict CVD^4^ and is seen as an important marker of HTN related asymptomatic organ damage^4^. PWV has been postulated to be a part of the pathophysiologic mechanisms that lead to HTN and it has been shown to predict the onset of HTN in longitudinal studies^5-7^.

PWV and HTN are closely related in pathophysiology, with chronic high blood pressure leading to arterial wall damage through various mechanisms like mechanical stress, endothelial dysfunction, and inflammation ^11^. African Americans (AA) were found to have stiffer carotid arteries compared to European American, indicating a potential difference in the progression of arterial stiffness between the two ethnic groups ^12^. The authors of the study suggested that large artery stiffening may occur earlier or be more accelerated in African Americans, possibly due to earlier exposure to multiple risk factors.

HTN is a highly heritable trait and various large-scale genome wide association studies (GWAS) have identified many associated novel and well validated single nucleotide polymorphisms^15-17^. Although less studied from a genetic standpoint, GWAS have also provided evidence for the genetic basis of arterial stiffness^18,19^. However, very little is known about the role of lncRNA in arterial stiffness and its subsequent impact on HTN. Research indicates that the gene expression patterns in peripheral blood cells mirror those in the vascular wall ^20^. This finding suggests that analyzing blood gene expression profiles could be a valuable method for investigating the molecular mechanisms underlying arterial disease ^21^. However, very few studies explored the link between arterial stiffness and gene expression in peripheral blood. Two of such studies utilized real-time reverse transcription polymerase chain reaction (qRT-PCR) to analyze a select group of genes ^22,23^. Recently a study by Logan et al. utilized mRNA sequencing (mRNA-seq) to compare transcriptome profiles in a modest sample size of twenty females with high and low arterial stiffness to understand early changes in arterial stiffness ^24^.

Our study aims to elucidate the transcriptomic footprint associated with arterial stiffness and its connection to hypertension in a large cohort of 615 African Americans. We utilize mRNA sequencing from whole blood samples to identify differentially expressed mRNAs between high and low arterial stiffness groups. Additionally, we explore the regulatory roles of long non-coding RNAs (lncRNAs) in modulating the mRNA expression. LncRNAs are transcripts longer than 200 nucleotides lacking protein-coding sequence that play an important role in cardiovascular disease pathophysiology ^25^ by influencing endothelial cell functions and vascular health ^26,27^. Our overarching goal is to enhance the understanding of how gene expression and lncRNAs relate to arterial stiffness and hypertension.

## MATERIAL AND METHODS

The data analyzed in this study originate from two primary sources: the GENomics, Environmental FactORs, and Social DEterminants of Cardiovascular Disease in African Americans STudy (GENE-FORECAST) and the Minority Health Genomics and Translational Research Bio-Repository Database (MH-GRID) project. Both studies received approval from the National Institutes of Health Institutional Review Board. They were conducted following local laws and institutional guidelines, with all participants providing written informed consent to participate.

### Phenotype Data

GENE-FORECAST is a comprehensive research platform designed to utilize a multi-omics systems biology approach for detailed analysis of minority health and disease in African Americans. It established a cohort of self-identified U.S.-born African American men and women (ages 21-65) from the Washington D.C. area. The data were randomly divided into two equal groups, each consisting of 218 samples, to create discovery and replication datasets. This approach allowed for the analyses to be conducted in the discovery dataset and subsequently validated in the replication dataset. The baseline characteristics of the 436 samples with PWV measurements included in the analysis are presented in Table 1. Hypertension status was determined using the mean of three blood pressure (BP) measurements and consideration of antihypertensive medication. Subjects were classified as hypertensive if they had a systolic blood pressure (SBP) ≥ 140 mmHg, a diastolic blood pressure (DBP) ≥ 90 mmHg, or were taking high BP medication. Conversely, subjects were classified as normotensive if they had optimal BP (SBP ≤ 120 mmHg and DBP ≤ 80 mmHg) without the use of BP medication. Notably, 60% of hypertensive subjects in the study were not on any BP medication, which accounts for the elevated SBP and DBP values observed in Table 1.

**Table 1:** Baseline characteristics of the GENE-FORECAST samples in the discovery and replication datasets used to assess the relationship between mRNA and PWV and between.

| Characteristics | Discovery (n = 218) Mean or Count SD or Proportion |  | Replication (n = 218) Mean or Count SD or Proportion |  |
| --- | --- | --- | --- | --- |
| Pulse Wave Velocity (m/s) | 7.54 | 1.72 | 7.62 | 1.7 |
| Age (years) | 49 | 12 | 47 | 12 |
| **Sex** |  |  |  |  |
| Female | 156 | 71.60% | 153 | 70.20% |
| Male | 62 | 28.40% | 65 | 29.80% |
| **Hypertension** |  |  |  |  |
| Normotensive | 46 | 21.10% | 43 | 19.70% |
| Hypertensive | 172 | 78.90% | 175 | 80.30% |
| **Systolic Blood Pressure (mmHg)** | 166 | 37 | 165 | 36 |
| **Diastolic Blood Pressure (mmHg)** | 75 | 10 | 76 | 10 |
| **Body Mass Index (kg/m<sup>2</sup>)** | 31 | 7 | 32 | 6 |

MH-GRID is a study of hypertension among self-identified African Americans aged 30 to 55. The data for this analysis are from a sub-study involving 179 participants at the Morehouse School of Medicine in Atlanta, Georgia. PWV was not measured in MH-GRID; these data were used to provide an independent validation of the gene expression-hypertension relationship observed in the GENE-FORECAST cohort. In the MH-GRID study, hypertension was based on high blood pressure (BP) medication; the data includes 78 hypertensives (individuals on at least two blood pressure medications, including one diuretic) and 101 normotensives (individuals with optimal blood pressure, systolic/diastolic BP ≤ 120/80, without medication).

### Pulse Wave Velocity

PWV was measured using the SphygmoCor XCEL (AtCor Medical) device as previously described ^28,29^. Briefly, PWV was determined by recording the pulse waves of the carotid and femoral arteries, and calculating the ratio of the distance between the pulse measuring sites to the time delay between the carotid and femoral pulse waves. The distance was measured with a non-stretchable tape from the suprasternal notch to the carotid site and from the suprasternal notch to the femoral site. The former distance was subtracted from the latter and used in the calculation of PWV.

### Transcriptome Data

The transcriptome data include messenger RNA sequencing from whole blood samples. Total RNA was extracted using the MagMAXTM for Stabilized Blood Tubes RNA Isolation Kit according to the manufacturer’s protocol (Life Technologies, Carlsbad, CA). For library preparation, total RNA samples were transformed into indexed cDNA sequencing libraries with Illumina’s TrueSeq kits, and ribosomal RNA (rRNA) was removed.

Sequencing for the GENE-FORECAST samples involved paired-end sequencing on the Illumina HiSeq2500 and HiSeq4000 platforms, achieving a minimum sequencing depth of 50 million reads per sample. Similarly, the MH-GRID samples underwent paired-end sequencing on the Illumina HiSeq2000 platform (Illumina, USA), also with a minimum depth of 50 million reads per sample.

mRNA expression was quantified using a bioinformatics pipeline from the Broad Institute, used by the Genotype-Tissue Expression (GTEx) project, with the pipeline details available on GitHub ^30^. Transcripts not reaching an expression threshold of 2 counts per million (CPM) in at least 3 samples were excluded. The expression data were normalized using the Trimmed Mean of M-values (TMM) method ^31^, which is optimal for read count data. Principal component analysis (PCA) was performed to detect and exclude sample and transcript outliers. After applying these quality control filters, 17,947 protein-coding mRNAs and 9,645 lncRNAs were retained for subsequent statistical analyses.

## STATISTICAL ANALYSES

The differential expression analysis focused on the extremes of the PWV distribution, specifically the upper (76 samples) and lower tertiles (66 samples), because the tails of the distribution can help identify mRNA and lncRNA transcripts and pathways that are strongly associated with very high or very low PWV, which might be more pronounced and biologically relevant than associations found when examining the entire range PWV. Furthermore, focusing on the tails of the distribution can increase the statistical power to detect differential expression because extreme groups are more likely to show clear differences, reducing the noise and variability that might obscure significant findings in our heterogeneous study population.

In the initial step, the R library *edgeR* was utilized to identify transcripts differentially expressed between the upper and lower tertiles of PWV, among the 17,947 mRNA and 9,645 lncRNAs that passed quality control filtering. *edgeR* fits which a negative binomial model to the transcripts read counts (i.e. expression) and computes likelihood ratio tests for the coefficients in the model. The model was adjusted for age and sex in both the discovery dataset and the replication dataset, both from GENE-FORECAST. Transcripts were considered significantly differentially expressed (DE) if the Benjamini and Hochberg (BH) ^32^ false discovery rate (FDR) adjusted p-value ≤ 0.05 in the discovery dataset, the nominal p-value ≤ 0.05 in the replication dataset and the log fold change (logFC) of the difference, between the upper and lower tertile group, is in the same direction.

Subsequently, the relationship between differentially expressed lncRNAs and differentially expressed mRNAs was investigated. An adapted version of the R library *MatrixEQTL* was utilized to identify associated lncRNA-mRNA pairs. The library was used to fit a linear regression model, where the expression level of each mRNA is regressed on the lncRNA expression level. The model was adjusted for age and sex and the p-values of the associations were adjusted for multiple testing by the BH approach. A lncRNA-mRNA association was deemed statistically significant and replicated if the multiple testing-adjusted p-value ≤ 0.05 in the GENE-FORECAST discovery dataset and the nominal p-value ≤ 0.05 in the replication GENE-FORECAST dataset, with consistent direction of the beta value in both datasets.

Finally, the relationship between hypertension status and the mRNAs differentially expressed by PWV and associated with lncRNAs was evaluated in the whole GENE-FORECAST sample set (347 hypertensives vs. 89 normotensives) and validated in MH-GRID (78 hypertensives vs. 101 normotensives). The model was adjusted for age and sex in both datasets and mRNA transcripts were considered significantly differentially expressed if the BH FDR adjusted p-value ≤ 0.05 in the GENE-FORECAST, the nominal p-value ≤ 0.05 in the MH-GRID dataset with consistent direction of the logFC in both datasets.

## RESULTS

### mRNAs and lncRNAs associated with PWV

Differential expression (DE) analysis was conducted to identify mRNAs and lncRNA differentially expressed between individuals in upper vs. lower tertile of PWV, in the GENE-FORECAST discovery and replication sets.

The mRNA differential expression (DE) analysis identified 1,035 mRNAs exhibiting statistically significant differential expression, characterized by an FDR-adjusted p-value ≤ 0.05 in the discovery dataset and a p-value ≤ 0.05 in the replication dataset. A comprehensive list of these differentially expressed mRNAs, along with their respective log-fold change (logFC) values and corresponding raw and adjusted p-values, is provided in the Supplemental Material (Table T1). Table 2 provides a summary of the number of lncRNAs associated with each of the 22 mRNAs reported in Figure 2 shows a graphical representation of this differential expression, highlighting the 22 mRNAs of particular interest in the subsequent analyses. All but 3 of the mRNA are up-regulated in the group with high PWV.

**Table 2:** Number of lncRNAs associated with each of the 22 differentially expressed mRNAs associated with lncRNAs and linked to hypertension in subsequent analyses.

| mRNA | Name | # lncRNA |
| --- | --- | --- |
| *AHSP* | alpha hemoglobin stabilizing protein | 28 |
| *JCHAIN* | joining chain of multimeric IgA and IgM | 27 |
| *ARF3* | ADP ribosylation factor 3 | 30 |
| *CISH* | cytokine inducible SH2 containing protein | 30 |
| *TRIM14* | tripartite motif containing 14 | 30 |
| *CTDNEP1* | CTD nuclear envelope phosphatase 1 | 30 |
| *PLAGL2* | PLAG1 like zinc finger 2 | 30 |
| *TMEM106A* | transmembrane protein 106A | 31 |
| *GP1BA* | glycoprotein Ib platelet subunit alpha | 30 |
| *LY6E* | lymphocyte antigen 6 family member E | 28 |
| *CEBPB* | CCAAT enhancer binding protein beta | 30 |
| *ARHGAP17* | Rho GTPase activating protein 17 | 31 |
| *LGALS9* | galectin 9 | 29 |
| *JUP* | junction plakoglobin | 30 |
| *POLR2J2* | RNA polymerase II subunit J2 | 31 |
| *HIVEP3* | HIVEP zinc finger 3 | 30 |
| *VAMP2* | vesicle associated membrane protein 2 | 31 |
| *CHST12* | carbohydrate sulfotransferase 12 | 31 |
| *ENSG00000269242* | - | 31 |
| *SIGLEC14* | sialic acid binding Ig like lectin 14 | 30 |
| *UCP2* | uncoupling protein 2 | 30 |

**Figure 1.**
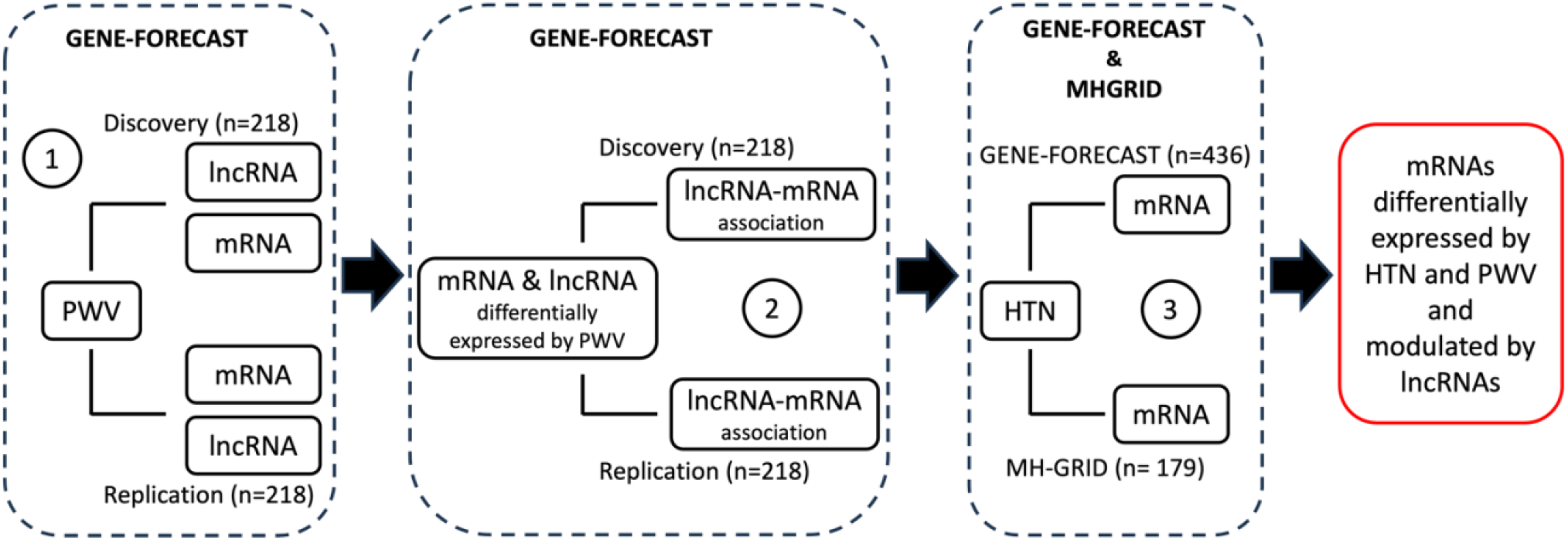
Summary of the analyses performed: (1) Identify and replicate mRNAs and lncRNAs differentially expressed by PWV, (2) identify and replicate lncRNA-mRNA pairs that are associated and (3) identify and validate mRNA differentially expressed by hypertension (HTN) status.

**Figure 2.**
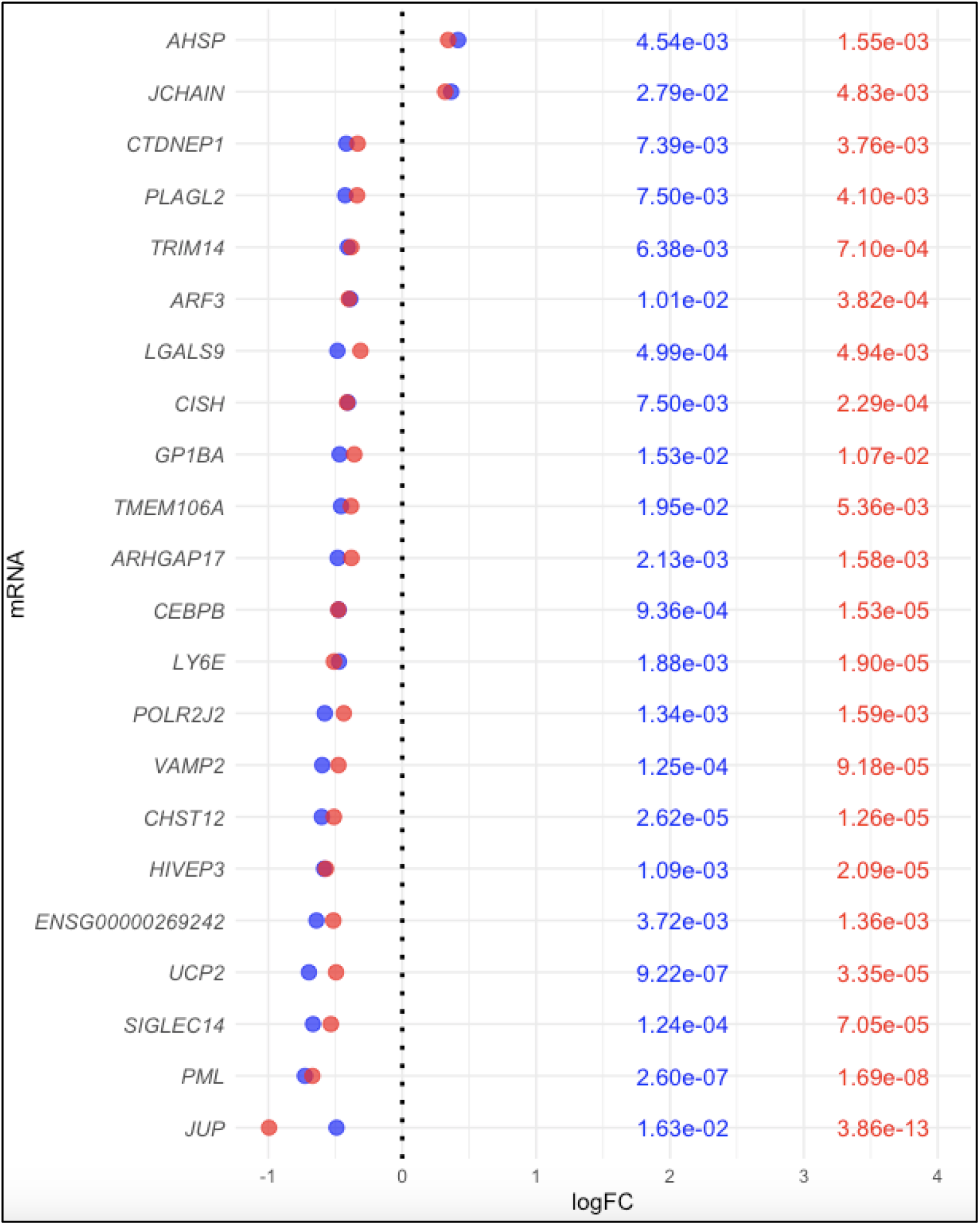
Summary of 22 of the mRNAs differentially expressed (DE) by PWV status in both datasets, including logFC and FDR-adjusted p-values in the discovery set (shown in blue) and p-values in the replication set (shown in red). The 22 mRNAs are also associated with lncRNA and DE by hypertension status in the subsequent analyses.

The lncRNA differential expression (DE) analysis identified 31 lncRNA exhibiting statistically significant differential expression, characterized by an FDR-adjusted p-value ≤ 0.05 in the discovery dataset and a p-value ≤ 0.05 in the replication dataset. More details about these differentially expressed lncRNAs, including their respective logFC values, corresponding raw and adjusted p-values, are provided in the Supplemental Material (Table T2). Figure 3 provides a graphical representation of this differential expression of the 31 lncRNAs, including 25 down-regulated and 6 up-regulated in the group with high PWV.

**Figure 3.**
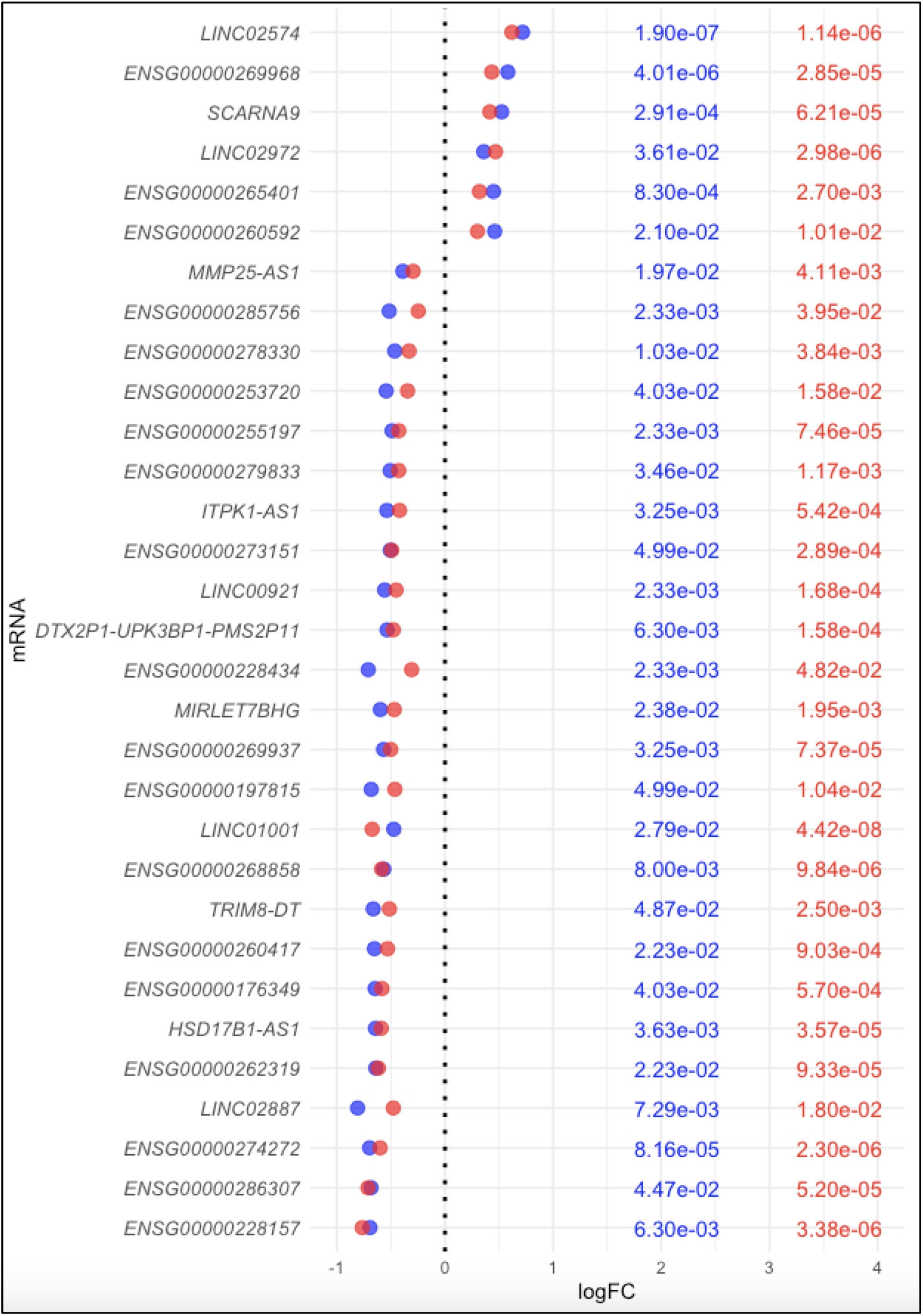
Summary of the 31 lncRNAs differentially expressed by PWV status in both datasets, including logFC and FDR-adjusted p-values in the discovery set (shown in blue) and p-values in the replication set (shown in red).

### lncRNA-mRNA pairs associated

Each of the protein-coding mRNA differentially expressed by PWV was tested for association with each of the 31 lncRNAs differentially expressed by PWV. The analysis identified 31,201 distinct lncRNA-mRNA associations statistically significant in discovery set (FDR adjusted p-value ≤ 0.05) and replication set (nominal p-value ≤ 0.05), with a beta coefficient in the same direction in both datasets. These associations involved all 31 lncRNAs, linked to the 1034 mRNAs. The detailed results of the lncRNA-mRNA association are provided in Supplemental Material (Table T3). Table 2 reports the number of lncRNAs associated with each of the 22 DE mRNAs shown in Figure 2.

### mRNAs associated with hypertension

The relationship between hypertension and the set of 1,034 mRNAs, differentially expressed by PWV and associated with lncRNAs, was evaluated to establish the connection between PWV and hypertension at the transcriptomic level. Differential expression analysis was performed to identify mRNAs exhibiting a significant difference of expression between normotensive and hypertensive subjects in the entire GENE-FORECAST dataset (comprising 89 normotensive and 347 hypertensive individuals) and the MH-GRID dataset (comprising 101 normotensive and 78 hypertensive individuals).

A total of 22 mRNAs were differentially expressed in the GENE-FORECAST dataset; these associations were subsequently validated in the MH-GRID dataset. All but one mRNA were up-regulated in hypertensive subjects. Figure 4 graphically presents the results of the analysis in both datasets.

**Figure 4.**
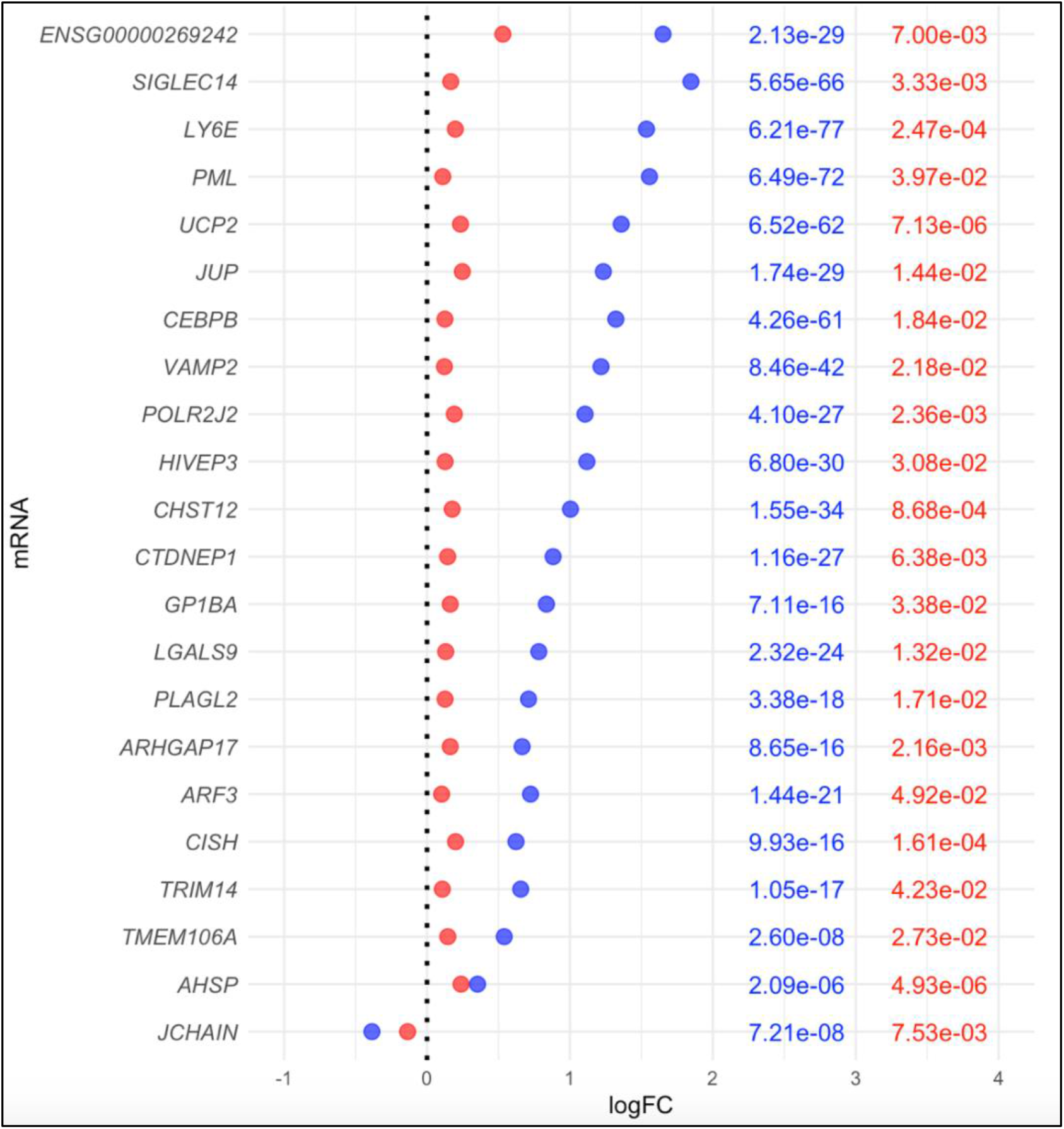
Summary of 22 of the mRNAs differentially expressed (DE) by hypertension and linked to PWV and lncRNAs in the two first analyses. The logFC and FDR-adjusted p-values in the GENE-FORECAST data are shown in blue whilst the logFC and p-values of the validation in MH-GRID are shown in red.

## DISCUSSION

### Summary

This study aimed to explore the transcriptomic footprint associated with arterial stiffness and its relationship to hypertension in 615 African Americans by identifying differentially expressed mRNAs and lncRNAs from whole blood samples.

The analyses revealed 22 mRNAs and 31 lncRNAs associated with arterial stiffness and hypertension across two independent datasets. The interaction between the lncRNAs and the mRNAs suggests that lncRNAs play a role in modulating gene expression related to pathways involved in arterial wall remodeling, which is an early indicator of chronically elevated blood pressure.

Among these findings, *UCP2* (Uncoupling Protein 2) emerged as a key gene of interest, prompting a focused discussion on its role in the context of arterial stiffness and hypertension.

### UCP2, Endothelial Function and Arterial Stiffness

UCP2 is a key mitochondrial antioxidant protein that plays a significant role in cardiovascular health by reducing oxidative stress, which is a major contributor to endothelial dysfunction and vascular stiffness ^33^. By uncoupling oxidative phosphorylation from ATP synthesis, UCP2 decreases the production of reactive oxygen species (ROS), mitigating cellular damage and improving endothelial function. In the context of arterial health, oxidative stress and mitochondrial dysfunction are pivotal factors contributing to endothelial dysfunction and vascular stiffness ^34^, which are essential in the pathogenesis of hypertension and atherosclerosis ^35^. By reducing oxidative stress, UCP2 helps protect against endothelial dysfunction and arterial stiffness, potentially influencing pulse wave velocity (PWV), a measure of arterial stiffness and an important predictor of cardiovascular risk ^36^.

### Mechanoregulation of UCP2 by Shear Stress

Mechanoregulation of *UCP2* by shear stress has been demonstrated, with unidirectional shear stress, statins, and resveratrol upregulating *UCP2* expression, whereas oscillatory shear stress and proinflammatory stimuli inhibit it through altered KLF2 (Krüppel-like factor 2) expression ^37^. KLF2 directly binds to the *UCP2* promoter, upregulating its transcription in endothelial cells ^38^. This regulation underscores the importance of hemodynamic forces in modulating *UCP2* expression and its downstream effects on vascular health ^39^. In animal models where *UCP2* expression is reduced or knocked out, there is an increase in mitochondrial oxidative stress, exacerbating conditions like hypertension and other cardiovascular disorders ^40^.

Intriguingly, in our analysis *UCP2* is down-regulated in subjects with high PWV (upper tertile) and up-regulated in hypertensives.

The downregulation of *UCP2* in subjects with higher PWV suggests that lower levels of *UCP2* may contribute to increased oxidative stress and endothelial dysfunction, leading to greater arterial stiffness. Whereas the upregulation of *UCP2* in hypertensive subjects might be a compensatory response to counteract the increased oxidative stress and inflammation associated with high blood pressure.

To further investigate this observation, we compared *UCP2* mRNA levels among normotensive individuals, specifically those in the upper tertile (n=30) versus the lower tertile (n=29) of PWV. We found that *UCP2* was significantly down-regulated in normotensives with high PWV (logFC = -0.46, p-value = 0.0007). This finding lends further credence to our hypothesis that downregulation of *UCP2* would lead to endothelial dysfunction, increased vascular stiffness and eventually hypertension. Furthermore, we compared *UCP2* mRNA levels between normotensive and hypertensive individuals within the upper tertile of PWV. *UCP2* was significantly up-regulated by about 2.5-fold in hypertensive individuals (logFC = 1.32, p-value = 3.80×10^−17^), lending further credence to our hypothesis that once hypertension develops there is an upregulation of *UCP2* as a compensatory mechanism to increase oxidative stress. These results suggest that by increasing *UCP2* levels, the body might attempt to mitigate the harmful effects of hypertension-induced oxidative stress.

### lncRNAs and *UCP2* Expression

The association between *UCP2* and various lncRNAs in our analysis reveals an intricate regulatory network. Most lncRNAs associated with *UCP2* expression were linked to its upregulation, indicating that these lncRNAs might enhance *UCP2* expression as a protective mechanism against oxidative stress and inflammation. For example, lncRNAs such as LINC02574, SCARNA9, LINC02972, MMP25-AS1, and ITPK1-AS1 have shown regulatory roles that could influence *UCP2* expression.

LINC02574 has been shown to inhibit viral replication and modulate immune responses, suggesting it might help mitigate vascular inflammation, a key factor in arterial stiffness ^41^. Vascular inflammation contributes to endothelial dysfunction, promoting the deposition of extracellular matrix components and the thickening of the arterial wall, which leads to increased arterial stiffness ^42^. By modulating immune responses, LINC02574 could play a role in maintaining vascular integrity and reducing stiffness.

SCARNA9 is involved in the modification of small nuclear RNAs (snRNAs) and the production of nucleolus-enriched fragments that form regulatory RNPs ^43^. These modifications affect cellular stress responses and metabolic processes within vascular cells, potentially influencing endothelial function ^44^. Proper endothelial function is essential for vascular elasticity and preventing stiffness ^45^. Therefore, SCARNA9’s role in stress response regulation could be crucial in maintaining healthy endothelial cells and vascular flexibility.

Furthermore, MMP25-AS1 regulates the expression of MMP25, reducing neutrophil infiltration and epithelial injury ^46^. Neutrophil infiltration into vascular tissues can exacerbate inflammation and promote atherosclerosis, contributing to arterial stiffness ^47^. By reducing neutrophil infiltration, MMP25-AS1 helps maintain vascular integrity and prevent arterial stiffness. This regulatory mechanism highlights the importance of MMP25-AS1 in controlling vascular inflammation and remodeling.

lncRNA ITPK1-AS1, identified in a competing endogenous RNA (ceRNA) network, modulates inflammation-related signaling pathways, thereby influencing vascular inflammation and stiffness ^48^. The ceRNA network involves interactions between lncRNAs, microRNAs (miRNAs), and mRNAs, forming a complex regulatory web that fine-tunes gene expression ^49^. By modulating inflammation-related signaling pathways, ITPK1-AS1 could influence the expression of genes involved in vascular remodeling and endothelial function, contributing to arterial stiffness ^50^.

### *UCP2* and Inflammatory Pathways

*UCP2* knockdown leads to the expression of genes involved in proinflammatory and profibrotic signaling, resulting in a proatherogenic endothelial phenotype ^51^. EC-specific Ucp2 deletion promotes atherogenesis and collagen production, while *UCP2* overexpression inhibits carotid atherosclerotic plaque formation in disturbed flow-enhanced atherosclerosis models ^52^. RNA-sequencing analysis revealed FoxO1 as a major proinflammatory transcriptional regulator activated by *UCP2* knockdown, and FoxO1 inhibition reduced vascular inflammation and atherosclerosis ^37,53^. *UCP2* is critical for the phosphorylation of AMPK (AMP-activated protein kinase), a key regulator of cellular energy homeostasis and a modulator of oxidative stress and inflammation. AMPK activation promotes endothelial NO synthase (eNOS) activity, enhancing NO production and improving endothelial function ^54^. Additionally, UCP2-induced inhibition of FoxO1 (forkhead box protein O1) reduces vascular inflammation and atherosclerosis. FoxO1 is a proinflammatory transcriptional regulator, and its inhibition by *UCP2* decreases the expression of inflammatory genes, contributing to reduced arterial stiffness ^55^.

The interplay between the lncRNAs and *UCP2* expression provides insights into the molecular mechanisms underlying the observed trends in PWV and hypertension. By enhancing *UCP2* expression, these lncRNAs likely play a protective role against oxidative stress and inflammation, crucial factors in the development of arterial stiffness. This regulatory network highlights the importance of lncRNAs in cardiovascular health and their potential as therapeutic targets for managing arterial stiffness and hypertension ^56^. For instance, the NF-κB pathway, a central mediator of inflammation, can be modulated by *UCP2. UCP2* reduces NF-κB activation by decreasing ROS levels, which are known to activate NF-κB. By reducing NF-κB activation, *UCP2* helps lower the expression of proinflammatory genes, thereby preventing endothelial dysfunction and arterial stiffness ^57^. The PI3K/Akt pathway is another critical signaling cascade involved in vascular health. This pathway promotes endothelial survival and function by enhancing NO production and reducing oxidative stress ^58^. *UCP2*, by reducing ROS levels, can enhance the activation of the PI3K/Akt pathway, thereby improving endothelial function and reducing arterial stiffness.

In addition to the specific roles of these lncRNAs, their regulatory impact on *UCP2* expression ties into broader inflammatory and oxidative stress pathways. Chronic inflammation and oxidative stress are key drivers of arterial stiffness, contributing to endothelial dysfunction, vascular remodeling, and reduced arterial compliance ^59^. By influencing *UCP2* expression, lncRNAs help modulate these processes, maintaining vascular health and preventing the progression of arterial stiffness.

The role of lncRNAs in cardiovascular health extends beyond their regulation of *UCP2*. For example, LINC02574 might not only inhibit viral replication but also reduce the inflammation caused by viral infections, which can exacerbate vascular stiffness ^41^. SCARNA9, through its involvement in snRNA modification and regulatory RNP formation, might influence the endothelial stress response and maintain cellular homeostasis, preventing endothelial dysfunction ^49^. MMP25-AS1, by regulating MMP25, reduces neutrophil-driven inflammation and extracellular matrix degradation, both of which contribute to vascular stiffening ^46^. ITPK1-AS1 and its role in the ceRNA network emphasize the complex interplay between various RNA molecules in regulating gene expression and maintaining vascular health ^49^.

### Hypertension-related Inflammation and Arterial Health

Hypertension-related inflammation plays a critical role in the pathogenesis and progression of arterial health issues, including endothelial dysfunction and arterial stiffness ^60^. In addition, inflammation can be triggered by the immune system, which can also raise blood pressure. For example, macrophages, neutrophils, and dendritic cells can contribute to hypertension by releasing inflammatory cytokines ^60^. Research suggests that chronic inflammation plays a role in the development and maintenance of hypertension. Inflammation can damage blood vessel linings, which can lead to arterial stiffness and higher blood pressure ^61^.

Beyond UCP2, the other genes identified in this study are not known to have direct association with arterial wall biology or PWV. Nonetheless, several genes, including CISH, CEBPB, LGALS9, JUP, and SIGLEC14, are involved in processes related to inflammation, immune regulation, and cell adhesion, which can indirectly affect vascular health.

For instance, inflammatory processes mediated by immune cells can lead to endothelial dysfunction and arterial stiffness ^60^; SIGLEC14 (sialic acid binding Ig like lectin 14) SIGLEC14, through its role in activating macrophages and neutrophils, may influence these processes by affecting the inflammatory state of the arterial wall. The SIGLEC family, including SIGLEC14, modulates immune responses through interactions with sialylated ligands ^62^. This modulation can affect the immune environment within the arterial wall, potentially impacting the progression of vascular disease.

## Conclusions

In summary, the regulatory roles of lncRNAs in modulating UCP2 expression provide insights into the molecular mechanisms underlying arterial stiffness and hypertension. By regulating UCP2 expression, lncRNAs help modulate oxidative stress, inflammation, and endothelial function. This regulatory network underscores the importance of lncRNAs and UCP2 in maintaining vascular health and highlights potential therapeutic targets for managing arterial stiffness and hypertension.

Moreover, several genes, including CISH, CEBPB, LGALS9, JUP, and SIGLEC14, were implicated in inflammation and immune regulation, indirectly affecting vascular health. These insights advance our understanding of the molecular mechanisms linking gene expression to arterial stiffness and hypertension. Overall, our findings underscore the importance of lncRNA-mediated regulation in cardiovascular health and highlight potential targets for therapeutic interventions.

The interactions between lncRNAs and mRNAs, particularly involving UCP2, highlight their roles in modulating gene expression related to arterial wall remodeling and oxidative stress management. By linking gene expression to arterial stiffness and hypertension, the study contributes to the understanding of cardiovascular health in African Americans, potentially guiding future research.

### Strengths and limitations

The study benefits from a large cohort of 615 African Americans, enhancing the statistical power and generalizability of the findings. Utilizing mRNA sequencing from whole blood samples, it provides a comprehensive analysis of the transcriptomic footprint associated with arterial stiffness and hypertension. Both discovery and validation datasets (GENE-FORECAST and MH-GRID) report a strong relationship between hypertension status and the expression of mRNA modulated by lncRNA, adding robustness to the findings. The identification of significant lncRNA-mRNA associations provides valuable insights into the regulatory mechanisms underlying arterial stiffness and hypertension.

A significant limitation is that pulse wave velocity was not measured in the MH-GRID dataset, preventing the validation of the transcriptome analyses related to PWV in this cohort. However, by splitting the GENE-FORECAST into discovery and replication datasets, the analysis provides reliable results within a large transcriptome dataset. The study’s cross-sectional design limits the ability to infer causal relationships between gene expression and arterial stiffness or hypertension. The findings are specific to African Americans and may not be generalizable to other populations. The analyses conducted do not account for potential confounding factors such as diet, physical activity, and socioeconomic status, which could influence gene expression and cardiovascular health. And finally, using whole blood samples may not capture tissue-specific gene expression changes relevant to arterial stiffness and hypertension.

## Supporting information

Details of the results

## CONTRIBUTORS

AG designed the analysis. GG processed and conducted quality controls of the transcriptome and phenotype data. AG conducted the statistical analyses and led the project. AG, JDV and MA interpreted the results. AG, JDV, MA, and GG drafted and edited the manuscript. All authors reviewed and approved the final version of the manuscript.

## DATA SHARING STATEMENT

Additional information can be found in the Supplementary Material of this article. The data presented in this article cannot be publicly shared due to privacy restrictions. Requests to access the datasets should be directed to the corresponding author

## DECLARATION OF INTERESTS

The authors have no financial and/or personal relationships with other people or organizations that could inappropriately influence (bias) this work.

## ACKNOWLEDGEMENT

This research was supported by the Intramural Research Program of the National Human Genome Research Institute, National Institutes of Health. The author are grateful to Dr Gary Gibbons.

## DECLARATION OF GENERATIVE AI AND AI-ASSISTED TECHNOLOGIES IN THE WRITING PROCESS

During the preparation of this work the author(s) used chatGPT in order to check and correct language spelling and grammar. After using this tool/service, the author(s) reviewed and edited the content as needed and take(s) full responsibility for the content of the publication.

## Notes

### Competing Interest Statement

The authors have declared no competing interest.

